# Dysfunction of cAMP-PKA-calcium signaling axis in striatal medium spiny neurons: a role in schizophrenia and Huntington’s disease pathogenesis

**DOI:** 10.1101/2021.10.08.463614

**Authors:** Marija Fjodorova, Zoe Noakes, Daniel C. De La Fuente, Adam C. Errington, Meng Li

**Author notes:** Correspondence (M.F.), (M.L.).

## Abstract

**Background:** Striatal medium spiny neurons (MSNs) are preferentially lost in Huntington’s disease. Genomic studies also implicate a direct role for MSNs in schizophrenia, a psychiatric disorder known to involve cortical neuron dysfunction. It remains unknown whether the two diseases share similar MSN pathogenesis or if neuronal deficits can be attributed to cell type-dependent biological pathways. Transcription factor BCL11B, which is expressed by all MSNs and deep layer cortical neurons, was recently proposed to drive selective neurodegeneration in Huntington’s disease and identified as a candidate risk gene in schizophrenia.

**Methods:** Using human stem cell-derived neurons lacking BCL11B as a model, we investigated cellular pathology in MSNs and cortical neurons in the context of these disorders. Integrative analyses between differentially expressed transcripts and published GWAS datasets identified cell type-specific disease-related phenotypes.

**Results:** We uncover a role for BCL11B in calcium homeostasis in both neuronal types, while deficits in mitochondrial function and protein kinase A (PKA)-dependent calcium transients are detected only in MSNs. Moreover, BCL11B-deficient MSNs display abnormal responses to glutamate and fail to integrate dopaminergic and glutamatergic stimulation, a key feature of striatal neurons *in vivo*. Gene enrichment analysis reveals overrepresentation of disorder risk genes among BCL11B-regulated pathways, primarily relating to cAMP-PKA-calcium signaling axis and synaptic signaling.

**Conclusions:** Our study indicates that Huntington’s disease and schizophrenia are likely to share neuronal pathogenesis where dysregulation of intracellular calcium levels is found in both striatal and cortical neurons. In contrast, reduction in PKA signaling and abnormal dopamine/glutamate receptor signaling is largely specific to MSNs.

## Introduction

Inhibitory γ-amino butyric acid (GABA)-releasing medium spiny neurons (MSNs) are the principal projection neurons of the basal ganglia, receiving inputs from both cortical glutamatergic neurons and midbrain dopaminergic neurons (mDA). MSNs are critically involved in a variety of essential functions including voluntary motor control, habit learning and reward processing, and dysfunction followed by loss of this neuron population underlies Huntington’s disease (HD). Growing body of evidence identifies loss of function of a transcription factor B-cell lymphoma/leukemia 11B (BCL11B, also known as CTIP2) as a driving force behind selective neuron degeneration in HD.

*BCL11B* is expressed by all MSNs and is required for transcriptional regulation of striatal genes, patch-matrix organization, spatial learning and working memory (1–4). *BCL11B* is also highly expressed by cortical layer V/VI neurons where it plays a role in corticospinal motor neuron fate specification and axon development (5, 6). BCL11B protein level is reduced in both human and rodent mutant huntingtin (mHTT)-expressing cells resulting in mitochondrial deficits prior to the onset of MSN death (7–10). We have recently demonstrated that human BCL11B-deficient and HD MSNs share dysregulated gene expression as well as present with deficits in signature striatal protein phosphorylation, DARPP32 and GLUR1 (11). In addition to the striatum, *BCL11B*-expressing neurons in other brain regions are affected in the later stages of HD, such as the cortex, hippocampus, and the hypothalamus (8). Thus, these findings point to an important role for BCL11B in HD pathogenesis.

Loss of function mutations in the *BCL11B* gene have also been identified to cause immunodeficiency and neurodevelopmental delay with speech impairment and intellectual disability (12). Furthermore, several studies have recently demonstrated significant enrichment of polymorphisms increasing risk for schizophrenia (SCZ) in the *BCL11B* gene (13–15). It has been widely accepted that pathogenesis in neurodevelopmental and psychiatric disorders is driven by cortical interneuron and glutamatergic neuron dysfunction (16), but novel evidence suggests that MSNs also play an important distinct role. Several gene enrichment studies integrating single cell transcriptomics and large psychiatric GWAS datasets revealed that psychiatric risk variants were highly enriched in genes expressed by MSNs that differ from those genes expressed by cortical neurons (13, 17, 18). Considering that polymorphisms affect both pan-neuronal and subtype-specific neuronal genes, it is important and necessary to investigate whether resulting cellular pathology is distinct or shared between striatal and cortical cell populations. It also remains to be determined as to what extent HD and SCZ pathogenesis may overlap due to disrupted function of BCL11B and if the same or different biological pathways are at play in these conditions. Together these findings lead to the hypothesis that BCL11B regulates important signaling processes in MSN and cortical neurons, whether shared or distinct, that may be particularly vulnerable to psychiatric disorder risk variants and HD pathogenesis.

Using BCL11B-deficient human embryonic stem cells (hESCs) as a model to address the above questions, we demonstrate here a role for BCL11B in mitochondrial function, calcium signaling and dopamine/glutamate signal processing predominantly in MSNs with much milder deficits observed in cortical neurons. Our study reveals significant enrichment of psychiatric disorder risk genes in BCL11B-regulated signaling including cAMP-dependent protein kinase A (PKA) and DARPP32 signaling specifically in MSNs, and calcium and glutamatergic synaptic signaling in both neuronal types. Similar biological pathways are identified in HD suggesting a shared role for BCL11B in the pathogenesis of HD and SCZ. We therefore prioritize promising targets for further mechanistic investigations and development of new therapeutics in these conditions.

## Methods and Materials

### Cell culture and neuronal differentiation

All experiments were performed in HUES9 iCas9 hESC line and genome-edited derivatives lacking BCL11B protein (clones #4, #33, #34) (11). As in previous study, BCL11B knockout (BCL11B^KO^) and control lines were reliably differentiated into MSN, cortical glutamatergic and mDA neuronal subtypes using established protocols (**Figure 1A and S1**) (19–21).

**Figure 1.**
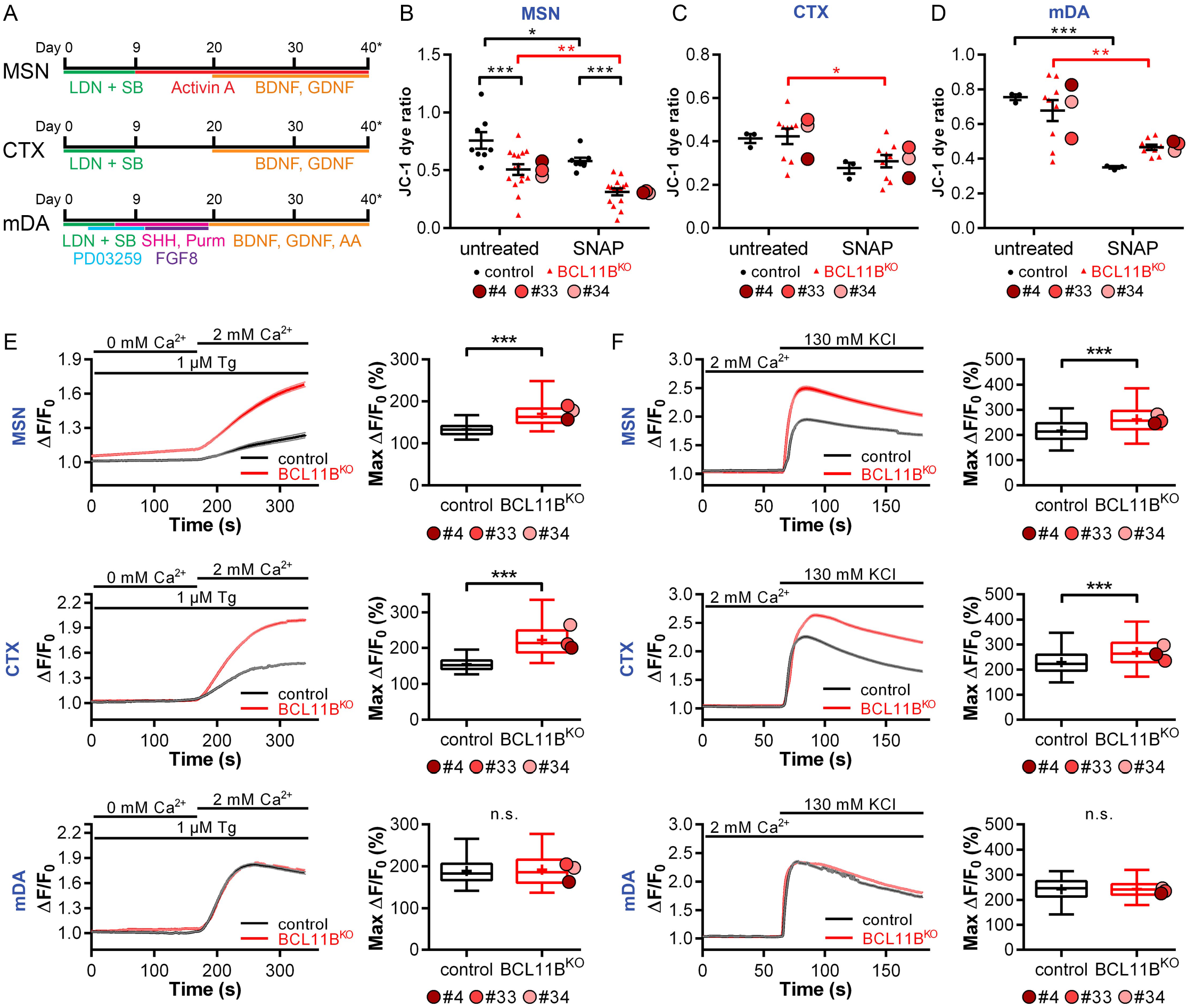
BCL11B^KO^ MSN and cortical neurons manifest signs of mitochondrial dysfunction and abnormal intracellular calcium regulation. (**A**) Schematic of MSN, CTX and mDA neuron differentiation protocols. Mitochondrial membrane potential as measured by JC-1 dye ratio at rest and after 24 hour treatment with 1000 μM SNAP in (**B**) MSNs, (**C**) CTX neurons and (**D**) mDA neurons at 40 DIV [two-way ANOVA with post hoc Bonferroni test: MSN – Genotype *F*_(1,42)_=30.552, *P*=1.9×10^-6^; Treatment *F*_(1,42)_=15.601, *P*=2.9×10^-4^; from left, in black: \*\*\**P*=4.6×10^-4^, 2.4×10^-4^; \**P*=0.024; in red: \*\**P*=0.001; CTX – Treatment *F*_(1,20)_=8.863, *P*=0.007; in red: \**P*=0.013; and mDA – Treatment *F*_(1,20)_=29.941, *P*=2.3×10^-5^; in black: \*\*\**P*=4.9×10^-4^; in red: \*\**P*=0.001], (**E**) Significantly increased store-operated channel-mediated Ca^2+^ entry in response to thapsigargin (Tg) is observed in BCL11B^KO^ versus control MSN and CTX neurons but not mDA neurons as demonstrated by Δ*F/F_0_* traces (left) and quantification of maximum Δ*F/F_0_* (right) [Mann-Whitney U test: MSN – *U*=63008.5, \*\*\**P*=2.4×10^-116^; CTX – *U*=33264, \*\*\**P*=5.8×10^-192^]. (**F**) Greater Ca^2+^ signals are induced in BCL11B^KO^ versus control MSN and CTX neurons in response to external KCI as demonstrated by Δ*F/F_0_* traces (left) and quantification of maximum Δ*F/F_0_* (right) [Mann-Whitney U test: MSN – *U*=515797, \*\*\**P*=3.3×10^-88^; CTX – *U*=344257, \*\*\**P*=1.5×10^-54^]. All line graphs and dot plots depict mean ± s.e.m. for each genotype. Box-and-whisker plots depict data for each genotype (center line – median,’+’ – mean, box limits – upper and lower quartiles, whiskers – 2.5 and 97.5 percentiles). Means for individual clones are indicated by red-shaded circles next to BCL11B^KO^ data; n.s., not significant; MSN, medium spiny neuron; CTX, cortical glutamatergic neuron; mDA, midbrain dopaminergic neuron. See also **Figures S1-2**.

### Mitochondrial Assay

The cell-permeant mitochondrial membrane potential sensor JC-1 (Thermo Fisher Scientific) was added directly to live cells at a final concentration of 1 μg/mL for 30 minutes at 37°C. Where indicated cells were pre-incubated with a nitric oxide donor (S)-Nitroso-N-acetylpenicillamine (SNAP, 1000 μM, Tocris) for 24 hours. Samples were analyzed on a flow cytometer according to the manufacturer’s protocol.

### Live-cell Calcium Imaging

Neuronal cultures were incubated with a Ca^2+^-sensitive probe Fluo-4-AM (Thermo Fisher Scientific) at a final concentration of 5 μM for 30 minutes at 37 °C. The solution was then replaced with pre-warmed artificial cerebrospinal fluid (aCSF) solution [2mM CaCl_2_, 142 mM NaCl, 2.5mM KCl, 1mM MgCl_2_, 10mM HEPES, 30mM D-glucose – all from Merck Sigma-Aldrich]. Neurons were imaged in a heated environmental chamber atop a Zeiss AG Axio Observer D1 inverted microscope stage. Where indicated aCSF was supplemented with one of the following: 1 μM thapsigargin, 10μM Roscovitine, 10 μM SKF-81297, 10 μM 8-bromo-cAMP (all from Tocris), 130 mM KCl, 100 μM DL-Glutamic acid (Merck Sigma-Aldrich). Additional procedure details can be found in the Supplement. Subsequent data processing was performed using FluoroSNNAP algorithm in MATLAB (MathWorks) according to the protocol (22).

### Electrophysiology

From 20 days of differentiation (DIV), MSN progenitors were cultured in astrocyte-conditioned medium supplemented with maturation factors according to an established protocol (23). Whole-cell patch-clamp recordings were acquired from control and BCL11B^KO^ #4 MSNs at 40 DIV in aCSF at room temperature. SKF81297 (10 μM) was added to aCSF where specified while DL-Glutamic acid (200 μM) was focally applied (30 ms) to the patched cell and the evoked current response recorded. Additional procedure details can be found in the Supplement. Data were analyzed with Clampfit software (Molecular Devices) then exported to and plotted using Origin (OriginLab).

### RNA Sequencing Data Analysis

Paired-end sequencing was performed at Oxford Genomic Centre on an Illumina HiSeq 4000 (Illumina, San Diego, USA) and RNA-seq data reported in this paper are available with the SRA accession numbers **PRJNA474679** and **PRJNA767962** (https://www.ncbi.nlm.nih.gov/sra/PRJNA474679 and https://www.ncbi.nlm.nih.gov/sra/PRJNA767962). The following R/Bioconductor packages and software were used to analyze differentially expressed genes and affected signaling pathways: DESeq2 (v1.14.1) (24), clusterProfiler (v3.12.14) (25) and Ingenuity Pathway Analysis (Qiagen).

## Results

### BCL11B-deficient MSNs display mitochondrial deficits

We showed previously that BCL11B^KO^ MSNs exhibited increased oxidative stress-dependent cell death (11). Since mitochondria are known to play an active role in this complex cascade of events in various models of neurodegenerative disorders (26), we set out to investigate whether loss of BCL11B would compromise mitochondrial health in MSN, cortical and mDA neurons. Mitochondrial membrane potential sensor JC-1 was used to measure metabolic activity, a key indicator of mitochondrial health (**Figure S2A**) (27). Mitochondria in BCL11B^KO^ MSNs were found to be significantly more depolarized than in control cells at 40 DIV, indicating a decreased metabolic activity in these neurons (**Figure 1B**).

To investigate whether this impairment in BCL11B^KO^ MSNs would render them more susceptible to oxidative stress, cells were treated with the nitric oxide donor SNAP, which was previously shown to generate reactive oxygen species (28). This induced a marked depolarization of mitochondrial membrane in both groups with a 2-fold greater amplitude in BCL11B^KO^ cultures (**Figure 1B** and **S2B**). We ascertained that the observed mitochondrial deficit was BCL11B-dependent in MSNs, by repeating this assay in striatal progenitors prior to BCL11B expression and detecting no deficit (**Figure S2C**).

We next investigated whether these mitochondrial deficits were specific to MSNs by performing the above experiment in cortical and mDA cultures. Although exposure to SNAP had a significant depolarizing effect in BCL11B^KO^ but not control cortical neurons after Bonferroni correction, there were no differences between the genotypes in either treatment group (**Figure 1C**). Also, no differences were detected due to loss of BCL11B in mDA neurons with both groups responding to SNAP similarly (**Figure 1D**). Taken together, our data suggests that BCL11B is required for regulation of mitochondrial membrane potential and has a protective role against oxidative stress, most prominently in MSNs and to a lesser extent in cortical neuron cultures.

### BCL11B^KO^ neurons present with abnormal intracellular calcium regulation

Mitochondrial dysfunction in neurons has previously been attributed to high intracellular calcium (Ca^2+^) levels, particularly in the context of neurological disorders (29). Given the impairments in mitochondrial health in BCL11B^KO^ neurons, we hypothesized that loss of BCL11B would result in deficits in intracellular Ca^2+^signaling and therefore performed calcium imaging in BCL11B^KO^ and control neuronal cultures at 50 DIV. To investigate the effect of BCL11B loss on the regulation of basal Ca^2+^ levels, intracellular Ca^2+^ was depleted using a Ca^2+^-free aCSF solution and then store-operated channel (SOC)-mediated Ca^2+^ entry was induced by application of the sarco/endoplasmic reticulum Ca^2+^ ATP-ase inhibitor, thapsigargin (30, 31). Significantly larger SOC-mediated Ca^2+^ signals, suggesting greater Ca^2+^ influx, were detected in both BCL11B^KO^ MSN and cortical but not mDA neurons compared to their respective controls (**Figure 1E**). Consistent with this finding, markedly larger Ca^2+^ signals in both BCL11B^KO^ MSN and cortical, but not mDA neurons, were also observed in response to depolarization of the plasma membrane by external potassium chloride (**Figure 1F**). This data indicates that BCL11B-dependent abnormal Ca^2+^ regulation and increased levels of intracellular Ca^2+^ may underlie the depolarized mitochondrial membrane potential and increased vulnerability to oxidative stress in MSN and cortical neurons described above.

### Signaling deficits in calcium transients are rescued by PKA activation in BCL11B^KO^ MSNs

Intracellular Ca^2+^ levels are also affected by Ca^2+^ influx and Ca^2+^ buffering during neuronal activity, which produces Ca^2+^ oscillations. Having established that BCL11B^KO^ MSN and cortical neurons exhibit a notable deficit in the regulation of intracellular Ca^2+^ levels, we next examined spontaneous activity evoked Ca^2+^transients (ΔCa^2+^) in these cells. In MSN cultures, smaller and less frequent ΔCa^2+^were detected in BCL11B^KO^ neurons compared to controls (**Figure 2A**). Indeed, this was reflected in a 36% decrease in spike amplitude and a 33% increase in interspike interval (ISI) in BCL11B^KO^ MSNs (**Figure 2B ‘MSN’**). Interestingly, this phenotype was specific to MSNs as no deficits were detected in BCL11B^KO^ cortical and mDA neurons either in ΔCa^2+^ amplitude or ISI (**Figure 2B ‘CTX’, ‘mDA’**).

**Figure 2.**
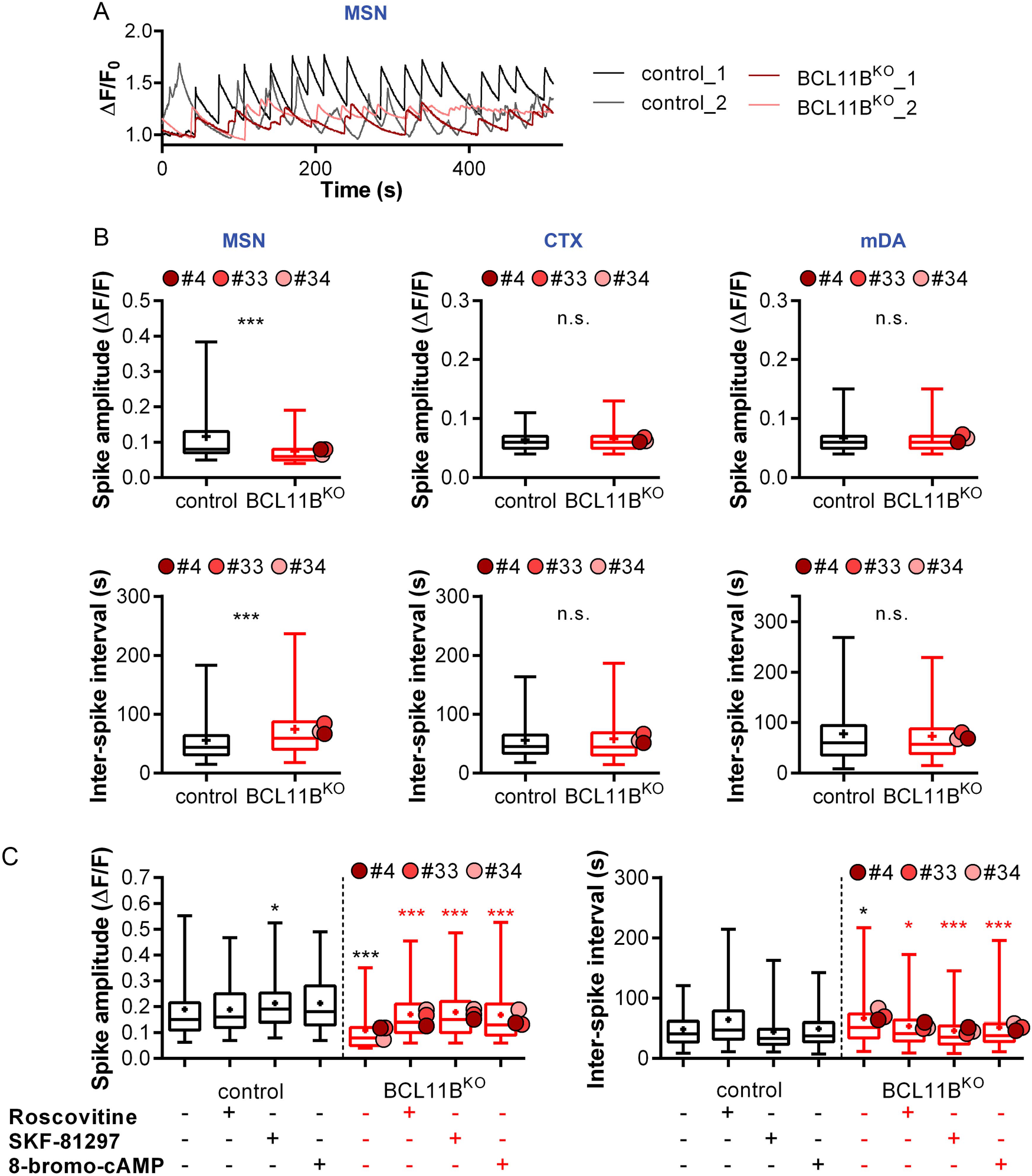
BCL11B knockout-induced abnormal calcium transient are rescued by PKA activation in MSNs. (**A**) Representative traces of spontaneous activity evoked Ca^2+^ transients in control and BCL11B^KO^ MSNs. (**B**) BCL11B-dependent reduced Ca^2+^ spike amplitudes and increased inter-spike intervals (ISIs) are present in MSN but not CTXor mDA neurons [Mann-Whitney U test: MSN amplitude – *U*=489091.5, \*\*\**P*=1.8×10^-85^; MSN interval – *U*=610109, \*\*\**P*=3.4×10^-31^]. (**C**) Lower Ca^2+^ spike amplitudes (left) and longer inter-spike intervals (right) in BCL11B^KO^versus control MSNs are rescued in a PKA-dependent manner with application of 10 μM Roscovitine, 10 μM SKF-81927 or 10 μM 8-bromo-cAMP [Kruskal-Wallis test with post hoc Bonferroni test: amplitude – X^2^_(7)_=275.877, *P*=8.5×10^-56^; from left, in black: \**P*=0.016, \*\*\**P*=3.1×10^-24^ vs untreated_control; in red: \*\*\**P*=2.5×10^-24^, 5.8×10^-25^, 3.7×10^-20^ vs untreated_BCL11B^KO^; ISI – X^2^_(7)_=78.147, *P*=3.3×10^-14^; from left, in black: \**P*=0.013 vs untreated_control; in red: \**P*=0.020, \*\*\**P*=3.8×10^-9^, 1.02×10^-4^ vs untreated_BCL11B^KO^], Box-and-whisker plots depict data for each genotype (center line – median,’+’ – mean, box limits – upper and lower quartiles, whiskers – 2.5 and 97.5 percentiles). Means for individual clones are indicated by red-shaded circles next to BCL11B^KO^ data; n.s., not significant; MSN, medium spiny neuron; CTX, cortical glutamatergic neuron; mDA, midbrain dopaminergic neuron.

We next sought to gain mechanistic insight into the reduction in spontaneous ΔCa^2+^in BCL11B^KO^ MSNs. Cross-talk between protein kinase A (PKA) and dopamine signaling pathways plays an important role in calcium signaling and DARPP32 phosphorylation in MSNs (32, 33). Given reduced levels of DARPP32-Thr34 phosphorylation and PKA signaling in BCL11B^KO^ MSNs identified previously (11), we hypothesized that activating PKA pathway would rescue ΔCa^2+^ deficits in these neurons. To this extent we compared the effects of drugs known to modulate PKA/DRD1-mediated calcium signaling in the striatum including CDK5 inhibitor Roscovitine, DRD1/DRD5 agonist SKF-81297 and cAMP analogue 8-bromo-cAMP (32, 34). All three treatments restored both the decreased ΔCa^2+^ amplitude and longer ISI in BCL11B^KO^ MSNs back to levels observed in controls (**Figure 2C**). Together, these findings suggest that loss of BCL11B disrupts PKA-dependent calcium signaling in MSNs, a deficit in spontaneous ΔCa^2+^ that is not observed in either cortical or mDA neurons.

### Abolished dopaminergic modulation of excitatory signaling and impaired glutamate-evoked responses in BCL11B^KO^ MSNs

Glutamatergic transmission in MSNs is known to be enhanced by dopamine acting on postsynaptic DRD1 (35). Considering abnormal dopaminergic and glutamatergic synaptic signaling in BCL11B-deficient MSNs suggested in prior study (11), we next inspected physiological implications of this by performing patch-clamp electrophysiology in MSN cultures at 40 DIV. While both control and BCL11B^KO^ neurons displayed similar basic membrane properties and fired action potentials with comparable rheobase, firing frequency was markedly higher in BCL11B^KO^ MSNs (**Figure 3A** and **S3A**). Glutamate-evoked currents were not significantly different between BCL11B^KO^ and control cells at baseline, however only control MSNs showed a significant increase in the current amplitude following treatment with DRD1/DRD5 agonist (**Figure 3B** and **S3B**).

**Figure 3.**
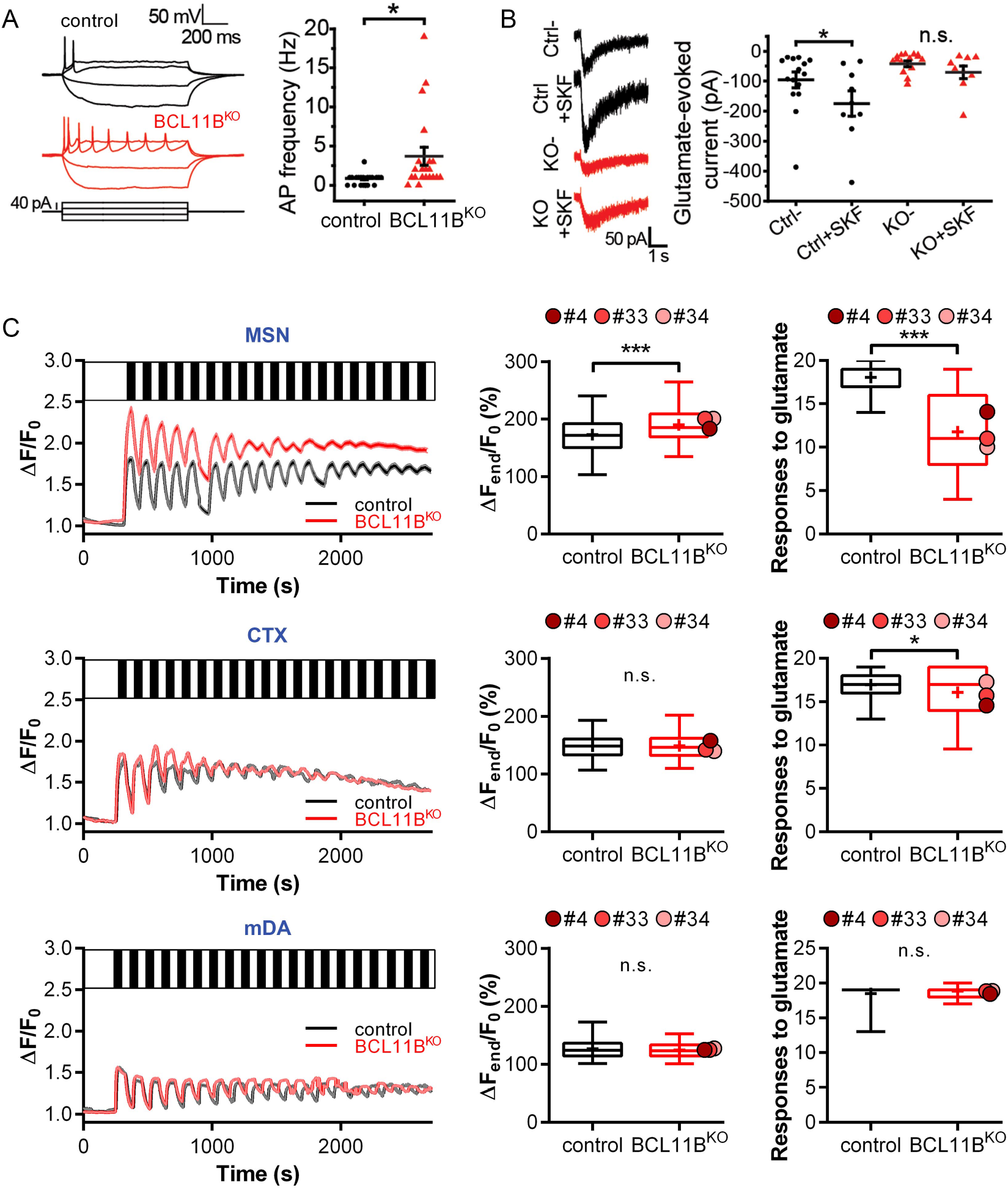
BCL11B^KO^ MSNs exhibit no dopaminergic modulation of excitatory signaling and present with impaired glutamate-evoked Ca^2+^ signals. (**A**) Representative traces of action potentials evoked by steps of current injection in control (black) and BCL11B^KO^ (red) MSNs (left). Increased firing frequency in response to current injection is observed in BCL11B^KO^ MSNs (right) [Mann-Whitney U test: *U*=90.5, \**P*=0.014], (**B**) Representative traces and quantification of current responses to glutamate (200 μM, 30 ms) in the absence or presence of SKF-81297 (10 μM) in control and BCL11B^KO^ MSNs. Only control MSNs show a significant enhancement of glutamatergic transmission induced by activation of dopaminergic signaling [two-way ANOVA with post hoc Bonferroni test: Genotype *F*_(1,40)_=8.860, *P*=0.005; Treatment *F*_(1,40)_=4.093, *P*=0.049; Ctrl- vs KO- *P*=0.123, Ctrl- vs Ctrl+SKF \**P*=0.038, KO- vs KÒ+SKF *P*=0.463, Ctrl+SKF vs KO+SKF \**P*=0.014], (**C**) Glutamate-evoked Ca^2+^ signals recorded in response to repetitive stimulation (100 μM, 1 minute, 19 pulses, black bars) in control and BCL11B^KO^ MSN, CTX and mDA neurons (left column). Glutamate-stimulated BCL11B^KO^ MSNs accumulate higher intracellular Ca^2+^ levels over time and respond to fewer glutamate pulses compared to control cells [Mann-Whitney U test: MSN Δ*F_end_/F_0_* – *U*=621047, \*\*\**P*= 1.8×10^-25^; responses – *U*=117472, \*\*\**P*=2.8×10^-153^]. Only a small deficit is present in the number of glutamate-evoked responses in BCL11B^KO^ CTX neurons, while no differences can be seen in mDA neurons [Mann-Whitney U test: CTX responses – *U*=287590, \**P*=0.043]. All line graphs and dot plots depict mean ± s.e.m. for each genotype. Box-and-whisker plots depict data for each genotype (center line – median,’+’ – mean, box limits – upper and lower quartiles, whiskers – 2.5 and 97.5 percentiles). Means for individual clones are indicated by red-shaded circles next to BCL11B^KO^ data; n.s., not significant; MSN, medium spiny neuron; CTX, cortical glutamatergic neuron; mDA, midbrain dopaminergic neuron. See also **Figure S3**.

Intracellular Ca^2+^ is delicately balanced and plays an indisputable role in determining neuronal excitability. Therefore, we next tested the effect of BCL11B loss on neuron response to excitatory stimulation by measuring ΔCa^2+^ evoked by glutamate pulses in MSN, cortical and mDA neurons (**Figure 3D**). BCL11B^KO^ MSNs initially exhibited larger glutamate-evoked ΔCa^2+^ but desensitized faster than control MSNs, a sign of excitotoxicity (**Figure 3D ‘MSN’**). Indeed, significantly greater increase in intracellular Ca^2+^ levels over time together with fewer glutamate-evoked responses were observed in BCL11B^KO^ MSNs compared to controls. Interestingly, this phenotype was mostly specific to MSNs as only a small deficit was detected in the number of glutamate-evoked responses in BCL11B^KO^ cortical neurons (**Figure 3D ‘CTX’**), while no differences were seen in mDA neurons due to loss of BCL11B (**Figure 3D ‘mDA’**).

A key measurement of neuronal functional maturity is the development of complex morphological features such as dendritic branching. Analysis of neuron morphology revealed no striking differences between control and BCL11B^KO^ MSNs (**Figure S3C-F**). In conclusion, although BCL11B-deficient MSNs exhibited enhanced intrinsic excitability, they presented with significantly impaired responsiveness to physiological stimuli, including glutamate-evoked Ca^2+^ signaling and DRD1-mediated modulation of glutamate-evoked currents, a feature characteristic of MSNs *in vivo* (35).

### cAMP-PKA-calcium signaling axis is driven by BCL11B-dependent transcription programs

To investigate molecular mechanisms and pathways leading to pathological changes in BCL11B^KO^ MSN and cortical neurons, we performed whole-transcriptome RNA-seq analysis at different stages of differentiation (**Table S1**). To elucidate specific biological processes regulated by BCL11B we performed KEGG and IPA pathway analysis of protein coding differentially expressed genes [DEGs; FDR(padj)<0.01]. Genes regulating calcium signaling, mitochondrial function and oxidative phosphorylation were found to be significantly altered in both neuronal types (**Figure 4A** and **Table S2**). Moreover, DA-DARPP32 feedback in cAMP signaling deficit was specific to MSN population while PKA and cAMP pathway dysregulation was stronger in MSNs but also present in cortical neurons. Furthermore, synaptic genes regulating dopamine, glutamate, and GABA neurotransmitters were significantly altered in MSNs from as early as 20 and 30 DIV. In contrast, only glutamatergic and GABAergic synapse signaling was affected in cortical neurons. In line with previously identified roles for BCL11B in the brain (3, 4), we also observed significant dysregulation of BDNF signaling specifically in MSNs, as well as abnormal learning and memory signaling in both neuronal types.

**Figure 4.**
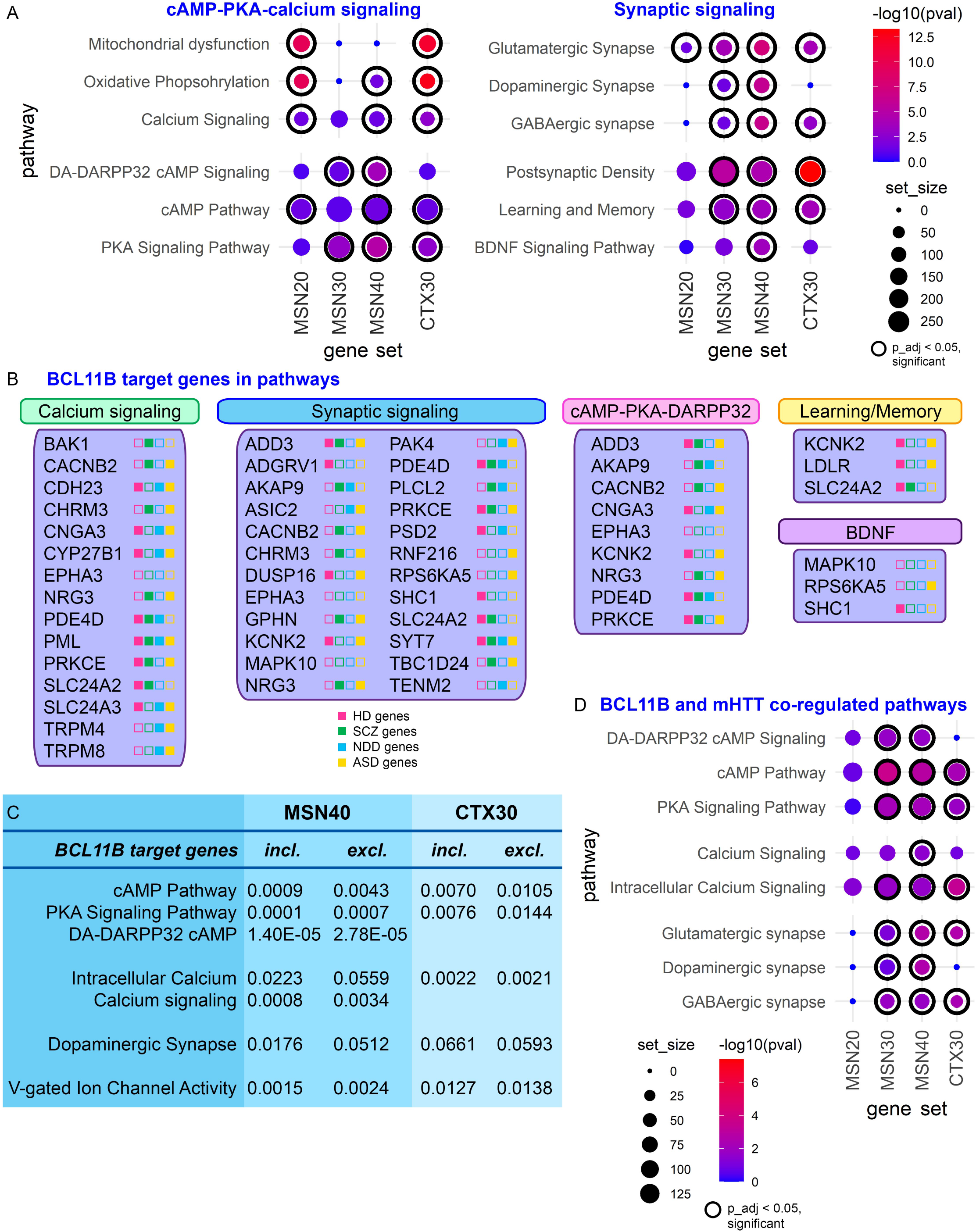
BCL11B-regulated transcription programs in MSN and cortical neurons and overlap with mHTT co-regulated pathways. (**A**) KEGG and Ingenuity Pathway Analysis (IPA) pathway analysis of DEGs at FDR(padj)<0.01 shows a significant enrichment of genes regulating cAMP-PKA-calcium signaling axis and synaptic signaling in both neuron types as well as DA-DARPP32 signaling specifically in MSNs (full gene set lists are presented in **Table S2**). (**B**) BCL11B target genes within shortlisted core biological processes affected by the loss of BCL11B in MSN and CTX neurons. Next to gene name, color icons indicate significant association (differential expression or identified risk variant) with Huntington’s disease (HD, pink), schizophrenia (SCZ, green), neurodevelopmental disorder (NDD, blue), and autism spectrum disorder (ASD, yellow). (**C**) Exclusion of BCL11B target genes from DEG list results in several pathways in MSNs becoming either drastically less significant or not significant at all, including cAMP-PKA-calcium signaling axis pathways and dopamine synapse signaling, while having almost no effect on signaling pathways in CTX neurons. (**D**) BCL11B-regulated signaling abnormalities in MSN and CTX neurons are shared by HD neurons (full gene set lists are presented in **Table S4**). In pathway graphs (A, D), dot size corresponds to gene set size, dot color corresponds to -log10(pval), where p_adj<0.05 dot is framed inside a black circle. MSN, medium spiny neuron; CTX, cortical glutamatergic neuron. See also **Figure S4 S5** and **Tables S1-4**.

To further elucidate the role of BCL11B in the identified pathogenic pathways, we asked whether the enriched pathways are driven by BCL11B target genes (**Figures 4B** and **S4**). Once BCL11B target genes were excluded from DEG list, several pathways in MSNs became either drastically less significant or not significant at all, including cAMP-PKA-calcium signaling axis pathways and dopamine synapse signaling (**Figure 4C** and **Table S2**). Interestingly, while BCL11B target genes appeared to play a weak role in cAMP-PKA signaling in cortical neurons, their exclusion did not affect calcium or dopamine synapse signaling in these cells. Transcriptomic data corroborates cellular pathologies identified in knockout neurons suggesting that BCL11B plays a role in mitochondrial and intracellular calcium signaling in both neuronal types, while its MSN-specific role converges on a few select pathways primarily regulating cAMP-PKA-calcium signaling axis and downstream glutamate/DA-DARPP32 neurotransmission.

### BCL11B-regulated signaling abnormalities are shared by HD neurons

BCL11B hypofunction has previously been demonstrated to play a role in MSN degeneration in HD by regulating mitochondrial signaling and protein phosphorylation (8, 11). We therefore investigated potential overlap between BCL11B- and mHTT-mediated transcriptomic changes by comparing our DEGs in both neuronal types with four publicly available datasets from human and mouse HD models (**Figure S5**) (7, 10, 36, 37). Gene set enrichment analysis indeed revealed that all MSN and cortical DEGs were significantly enriched for mHTT-regulated genes (**Table S3**). Further investigation of concordantly dysregulated genes between BCL11B^KO^ and HD models confirmed deficits in DA-DARPP32 signaling, cAMP-PKA-calcium signaling axis, synaptic signaling as well as oxidative phosphorylation and mitochondrial function predominantly in MSNs and to a lesser extent in cortical neurons (**Figure 4D** and **Table S4**). Interestingly, multiple BCL11B target genes in these pathways have been previously shown to be affected in HD (**Figure 4B**). These results provide further support for a regulatory role for BCL11B in cAMP-PKA-calcium signaling axis and downstream DA-DARPP32 neurotransmission events, processes that are particularly vulnerable to mHTT.

### A role for cAMP-PKA-calcium signaling axis in the pathogenesis of psychiatric disorders

A direct role for MSNs in psychiatric disease pathogenesis has been suggested by recent gene enrichment studies (13, 17, 18). Moreover, independent studies have identified and prioritized *BCL11B* gene among a few others as a candidate causal risk gene in SCZ (13–15). Considering these reports, we wonder whether the BCL11B-dependent neuronal phenotypes described in the present study contribute to the pathogenesis in psychiatric conditions. To gain support of this hypothesis we investigated BCL11B-regulated genes in MSN and cortical neurons, with particular interest in the signaling pathways responsible for the observed convergent and MSN-specific deficits, and their enrichment for risk variants in SCZ, neurodevelopmental disorder (NDD) and autism spectrum disorder (ASD). Neurological disease gene sets were collated by integrating findings from transcriptomics, GWAS datasets, and other functional genomics studies of SCZ (13, 18, 38–52), NDD (49, 51, 53) and ASD (42, 51, 54) (**Figure S5** and **Table S5)**. In line with neurodevelopmental component reported in SCZ and ASD, an overlap of about a third of genes was detected between these two conditions and NDD gene set.

Gene set enrichment analysis revealed that BCL11B^KO^ MSN DEGs were enriched for SCZ and NDD risk genes but not ASD risk genes at all stages of differentiation, while CTX30 DEGs were enriched for all disease risk genes (**Figure 5A** and **Table S6**). In MSN cultures, BCL11B-dependent enrichment of DA-DARPP32 and cAMP-PKA-calcium signaling functional terms was much stronger and appeared earlier for SCZ than NDD risk genes (**Figure 5B** and **Table S7**). Furthermore, SCZ risk variants were enriched in synaptic signaling affecting all three neurotransmitters while NDD risk genes were present only in glutamatergic and GABAergic synaptic pathways. In cortical cultures, dysregulated pathways were more enriched for SCZ and ASD than NDD risk variants. Interestingly, most BCL11B target genes in these pathways have been previously implicated in at least one of these three neurological disorders (**Figure 4B**). Together, these findings suggest that synaptic signaling and the cAMP-PKA-calcium signaling axis dysregulation may contribute to SCZ pathogenesis in the striatum. In cortical neurons in contrast, some of these BCL11B-dependent phenotypes, although affected less severely, may contribute to both SCZ and ASD pathogenesis.

**Figure 5.**
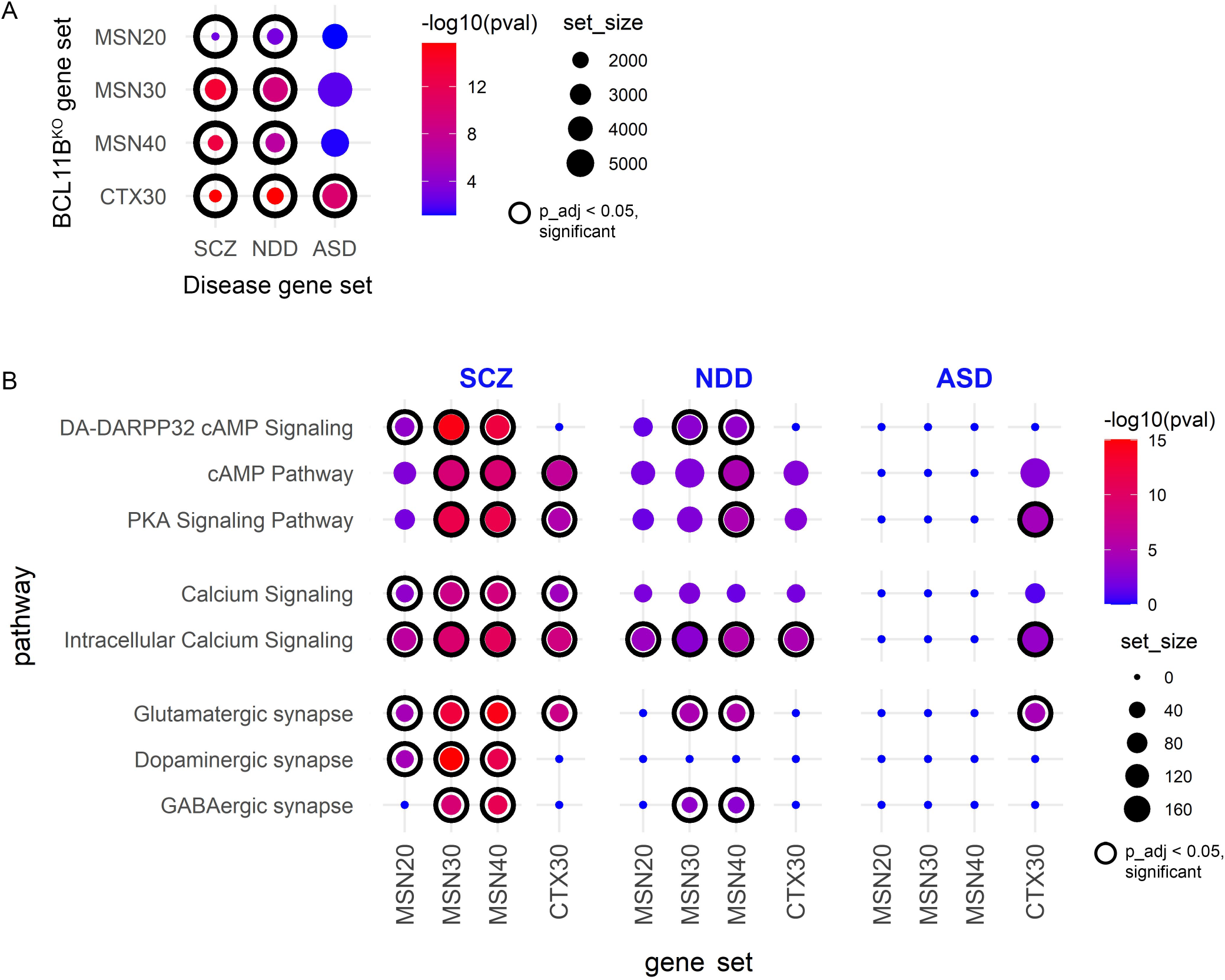
A role for BCL11B-dependent DA-DARPP32 and cAMP-PKA-calcium signaling axis in the pathogenesis of psychiatric disorders. (**A**) BCL11B-regulated genes in MSNs are significantly enriched for SCZ and NDD risk genes but not ASD risk genes at all stages of differentiation, while CTX30 DEGs were enriched for all disease risk genes. SCZ, schizophrenia; NDD, neurodevelopmental disorder; ASD, autism spectrum disorder. *P-values* were calculated from Fisher’s exact test followed by Bonferroni correction for multiple comparisons. (**B**) BCL1 IB-dependent signaling pathways in MSNs are predominantly enriched for SCZ but not NDD risk variants, while pathways in CTX were enriched for both SCZ and ASD risk genes (full gene set lists are presented in **Table S7**. In graphs dot size corresponds to gene set size, dot color corresponds to -log10(pval), where p_adj<0.05 dot is framed inside a black circle. MSN, medium spiny neuron; CTX, cortical glutamatergic neuron. See also **Figure S5** and **Tables S5-7**.

## Discussion

In this study, we provide evidence for a role for cortico-striatal transcription factor BCL11B predominantly in HD and SCZ than NDD and ASD pathogenesis. We further strengthen the hypothesis that MSN dysfunction contributes separately from cortical neuron pathology to psychiatric disease development. Such neuron subtype and multiple disorder cross-examination allows the opportunity to study gene expression patterns and determine ‘convergent’ versus ‘distinct’ BCL11B-regulated mechanisms contributing to different neurological disease pathogenesis in MSN and cortical neurons. Indeed, our gene enrichment analysis and functional assays reveals a limited number of core biological processes affected by the loss of BCL11B that are highly specific to MSN population such as cAMP-PKA-calcium signaling axis, DA-DARPP32 signaling, glutamate-evoked calcium signaling, and mitochondrial health (**Figure 6**). Intracellular calcium signaling deficit is the only phenotype common between MSN and cortical neurons, while other calcium and mitochondrial signaling deficits are only mildly present in cortical neurons. Transcriptomic analysis suggests that identified BCL11B-dependent biological processes are mainly concordant between HD and SCZ in MSNs, while cortical neuron pathways were also enriched for ASD risk variants. Furthermore, our study predicts involvement of BCL11B target genes in regulating these pathways, with many genes identified either as risk factors for or as being differentially expressed in psychiatric disorders.

**Figure 6.**
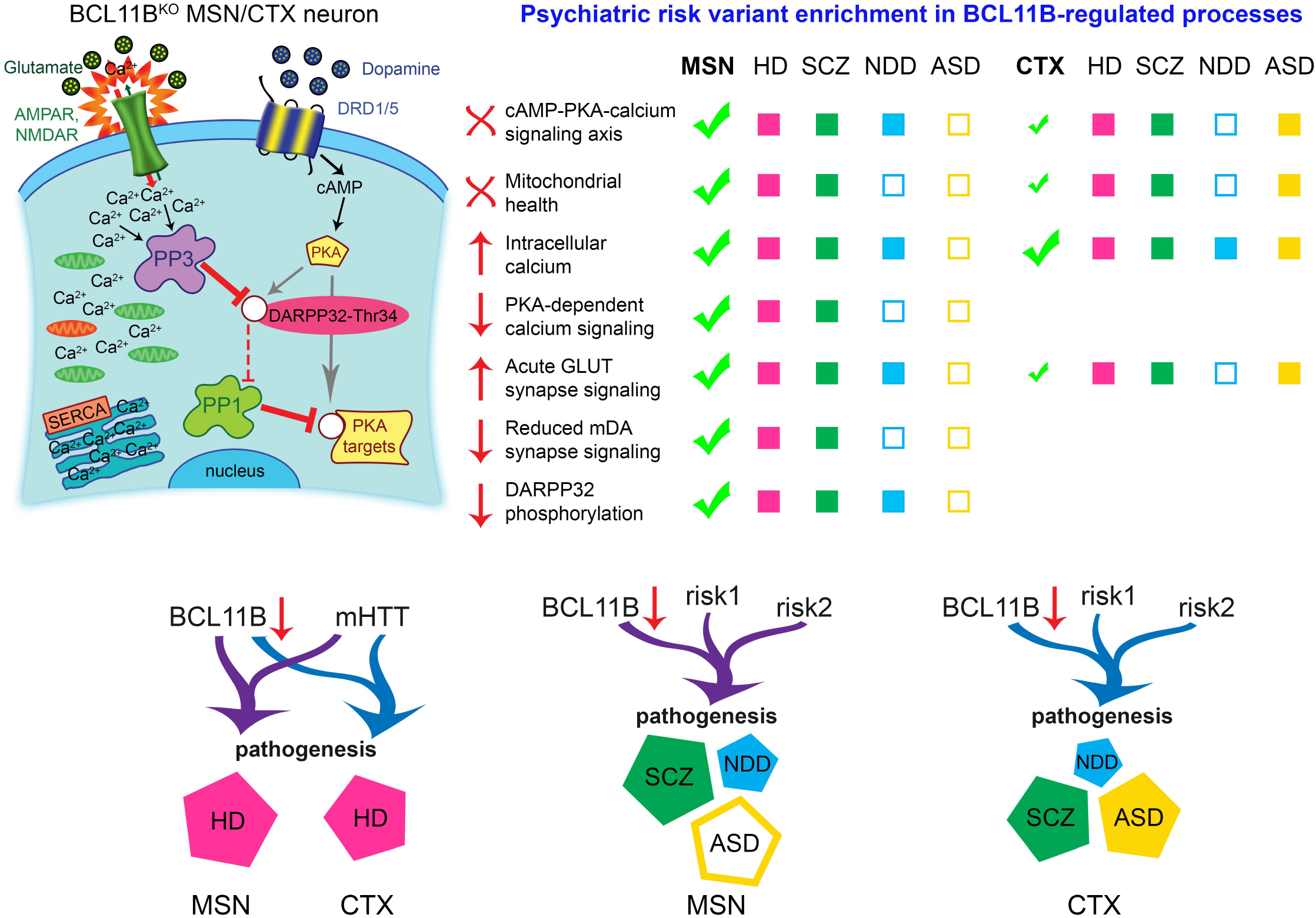
BCL11B-regulated molecular events in MSN and CTX neurons and their likely contribution to psychiatric disorder pathogenesis. Schematic of core biological processes affected by the loss of BCL11B summarizes phenotypes identified in current and previous study (left), including depolarized mitochondria (green), abnormal calcium levels, acute glutamatergic neurotransmission, disbalance in phosphatase levels, reduced cAMP-PKA signaling and downstream dephosphorylation of DARPP32 and other targets [adapted from (11)]. These core molecular changes are most prominent in MSNs compared to CTX neurons and show preferential enrichment for HD and SCZ genes suggesting that they are likely to contribute to the pathogenesis in these disorders (right). In CTX neurons in contrast, a fraction of these BCL11B-dependent phenotypes, although affected less severely, may also contribute to ASD pathogenesis. In summary, we propose that MSNs play a distinct role in psychiatric disease development compared to CTX neurons, where BCL11B hypofunction acts in tandem with mHTT in HD and other risk variants in psychiatric disorders to contribute to disease pathogenesis (bottom). MSN, medium spiny neuron; CTX, cortical glutamatergic neuron.

Interestingly, BCL11B target genes appear to drive cAMP-PKA-calcium signaling axis and dopamine receptor signaling pathways predominantly in MSNs. Most notable genes among them are *ADD3, CACNB2, CNGA3, KCNK2*, and *SYT7*, with significant downregulation of *ADD3* and *CNGA3* already confirmed in independent BCL11B^KO^ samples in our previous study (11). These five genes play a role in activation of cAMP-PKA-calcium signaling, synaptic signaling and excitation of neurons (55–59). Furthermore, they are either dysregulated in or considered to be risk factors for multiple neurological diseases including HD, SCZ, NDD, ASD, and bipolar disorder (7, 10, 36, 43, 44, 53, 60–64). Together, this evidence provides mechanistic insight into how BCL11B hypofunction may contribute to the disturbed cAMP-PKA-calcium and synaptic signaling in MSNs in HD and SCZ.

Intriguingly, transgenic expression of *Blc11b* in *STHdh^Q111^* HD cells (that exhibit reduced levels of Bcl11b) partially rescued mHTT-induced defects in mitochondrial metabolic activity (8). This suggests that restoring or increasing BCL11B levels can reverse at least some of the mHTT-driven impairments. Mitochondrial dysfunction and elevated intracellular Ca^2+^ levels are highly interdependent phenotypes in neurodegenerative diseases (29). Interestingly, like HD cells, BCL11B-deficient neurons present with mitochondrial deficits, vulnerability to oxidative stress and abnormal intracellular Ca^2+^ levels. While current results cannot definitively discriminate between increased Ca^2+^ influx and other mechanisms, such as impaired Ca^2+^ buffering, our data shows that regulation of basal Ca^2+^ levels is impaired by the loss of BCL11B and this could lead to intracellular Ca^2+^ overload; a proposed critical step in neurodegeneration (65). Moreover, same deficits in mitochondrial membrane potential as reported for BCL11B^KO^ MSNs here have been previously described in cortical neurons derived from SCZ patients (66). Although no deficits in either electrophysiological or spontaneous ΔCa^2+^ activity were found in cortical neurons in this study (66), reduced electrical activity in hippocampal CA3 neurons derived from SCZ patients has been reported elsewhere (67, 68), while no studies in patient MSNs have been conducted yet. We demonstrate that BCL11B-dependent oxidative phosphorylation as well as calcium signaling in MSNs are enriched for SCZ risk genes. This suggests that BCL11B-associated phenotypes may contribute to neuropathology in SCZ and that there might be yet undiscovered MSN dysfunction in this psychiatric disease development.

A connection between disturbed calcium signaling, excitatory stimulation and MSN apoptosis has been established in HD models (31, 69). We demonstrate that glutamate induces elevated Ca^2+^ responses in BCL11B^KO^ MSNs, a feature of excitotoxicity, which suggests that BCL11B hypofunction might be the cause of aberrant calcium signaling and MSN apoptosis observed in HD. Although BCL11B knockout appears to enhance intrinsic excitability of MSNs, it simultaneously impairs dopaminergic modulation of glutamate-mediated excitation. Together, these results provide strong evidence of a central role for BCL11B in regulating calcium homeostasis in MSNs and to some extent in cortical neurons, disruption of which in BCL11B-deficient and HD MSNs induces Ca^2+^ overload leading to excitotoxicity and pathological responses to physiological stimuli. Interestingly, BCL11B-associated glutamatergic and dopaminergic signaling phenotypes may also be present in MSNs in SCZ.

Furthermore, PKA signaling was previously implicated in modulating intracellular Ca^2+^ levels and Ca^2+^ oscillations (32, 70). In agreement with these findings, we demonstrate that deficits in spontaneous activity-evoked Ca^2+^ oscillations in BCL11B-deficient MSNs can be rescued in a PKA-dependent manner. Thus, we propose that BCL11B deficiency-driven disruption of calcium homeostasis in MSNs, together with dysregulated levels of protein phosphatases/kinases (11), result from reduced PKA signaling and lead to a marked decrease in phosphorylation of its targets. In addition, increased intracellular Ca^2+^ levels would induce overactivation of PP3-dependent DARPP32-Thr34 dephosphorylation (33). Identified disruption of PKA-regulated intracellular calcium signaling and phosphorylation of DARPP32 in MSNs has significant implications for many psychiatric and neurodegenerative diseases. These processes regulate transcriptional and behavioral responses of MSNs to pharmacological stimuli, including antidepressants, neuroleptics, and drugs of abuse (71). Interestingly, reduced levels of full-length DARPP32 and increased levels of DARPP32 isoforms lacking the crucial residue Thr34 were reported in SCZ patients (72, 73), which suggests disruption of DARPP32-Thr34 phosphorylation in SCZ MSNs. Moreover, abnormal postsynaptic PKA activity due to accelerated maturation of cortico-striatal circuits was demonstrated to cause behavioral abnormalities in *Shank3B^-/-^* mouse model of ASD (74). Indeed, we demonstrate that BCL11B-dependent cAMP-PKA-calcium signaling and DA-DARPP32 signaling pathways in MSNs are enriched for psychiatric disorder risk variants, pointing to a role for BCL11B-associated phenotypes in neuropathology in these disorders.

In conclusion, we identify BCL11B-regulated molecular mechanisms in striatal and cortical neurons and further strengthen the hypothesis that MSN dysfunction contributes separately from cortical neuron pathology to psychiatric disease development. We provide evidence that genetic susceptibility loci within BCL11B-regulated pathways may modulate a limited number of core disease-related biological processes in MSNs, including cAMP-PKA-calcium signaling, DA-DARPP32 and glutamate neurotransmission. We propose that BCL11B-associated phenotypes may contribute to neuropathology most significantly in HD and SCZ and identify modulation of PKA-dependent Ca^2+^ signals and protein phosphorylation as potential new therapeutic targets in the striatum.

## Acknowledgements

We thank Dr. Robert Andrews (Advanced Research Computing at Cardiff) for invaluable assistance with RNA-seq data analysis. RNA-seq was performed at the Oxford Genomics Centre. Thanks also to all members of the M.L. laboratory for helpful discussions during this study. This work was supported by EU framework program 7 *repair-HD* and UK Medical Research Council grants to M.L. and Jane Hodge Foundation Neuroscience Research Fellowship to A.C.E..

## Author Contributions

M.F. and M.L. conceived the study and designed experiments. M.F. carried out and analyzed the experiments. Z.N. performed and analyzed the electrophysiological studies with supervision from A.C.E.. D.C.F. contributed to the RNA-seq data analysis and interpretation. M.F. and M.L. wrote the manuscript.

