## Supplemental Information for "Dysfunction of cAMP-PKA-calcium signaling axis in striatal medium spiny neurons: a role in schizophrenia and Huntington’s disease pathogenesis"

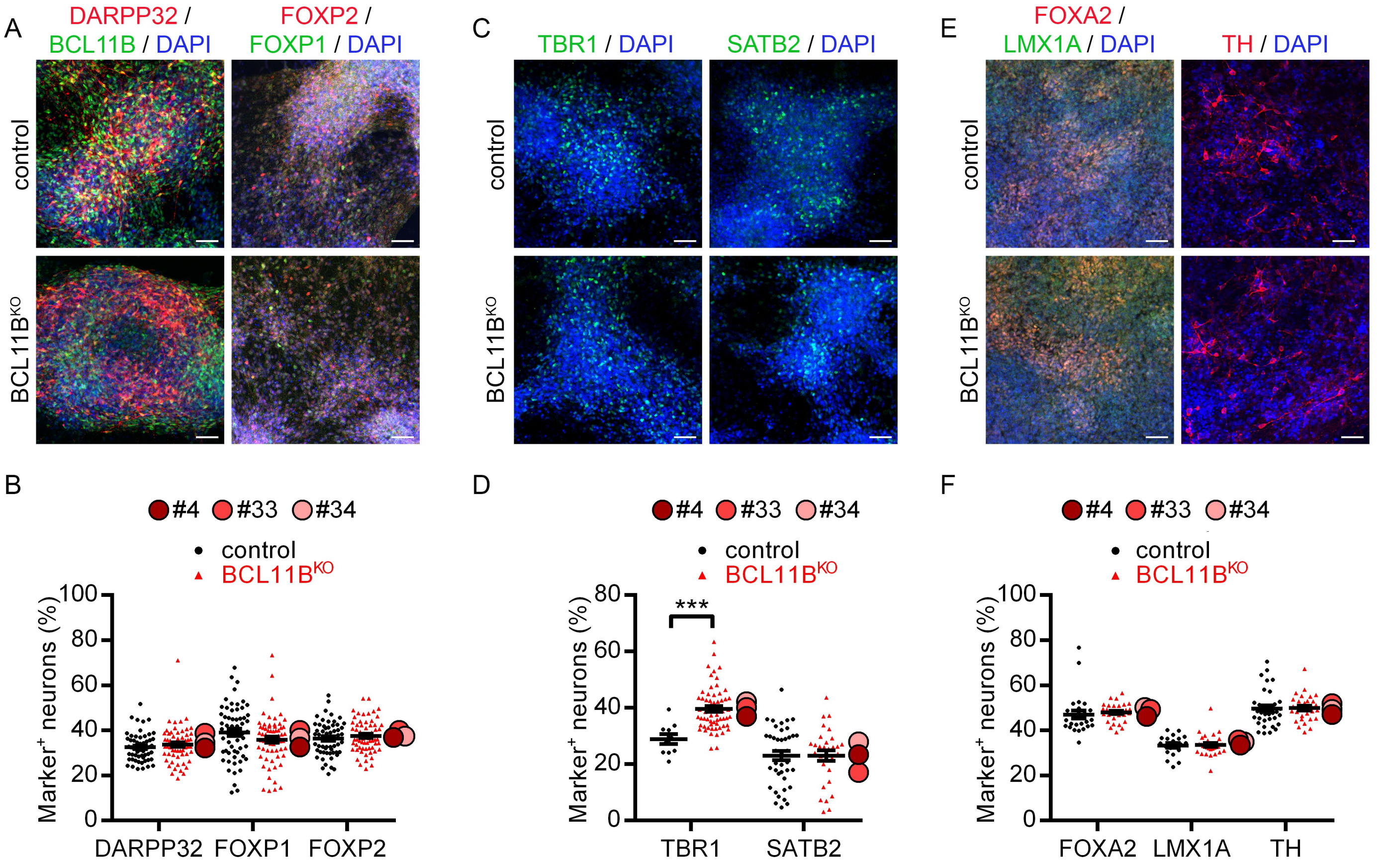


**Figure S1 (Related to Figure 1) – Loss of BCL11B has no effects on regional specification of hESC-derived MSN, cortical and mDA neurons.** Images (**A**) and quantification (**B**) of MSNs in control and BCL11B$^{KO}$ cultures at 40 DIV immunostained for BCL11B, DARPP32, FOXP1, FOXP2. The BCL11B antibody binds to exon 1 of the protein, which is upstream of the truncation. Images (**C**) and quantification (**D**) of cortical neurons in control and BCL11B$^{KO}$ cultures at 40 DIV immunostained for TBR1 and SATB2 [Mann-Whitney U test: TBR1 – $U$=66, *$P$=8.6 x10$^{-5}$]. Images (**E**) and quantification (**F**) of mDA neurons in control and BCL11B$^{KO}$ cultures at 40 DIV immunostained for FOXA2, LMX1A and TH. All scale bars: 50μm. All graphs show mean ± s.e.m. for each genotype, with the means for individual clones indicated by red-shaded circles next to BCL11B$^{KO}$ data.

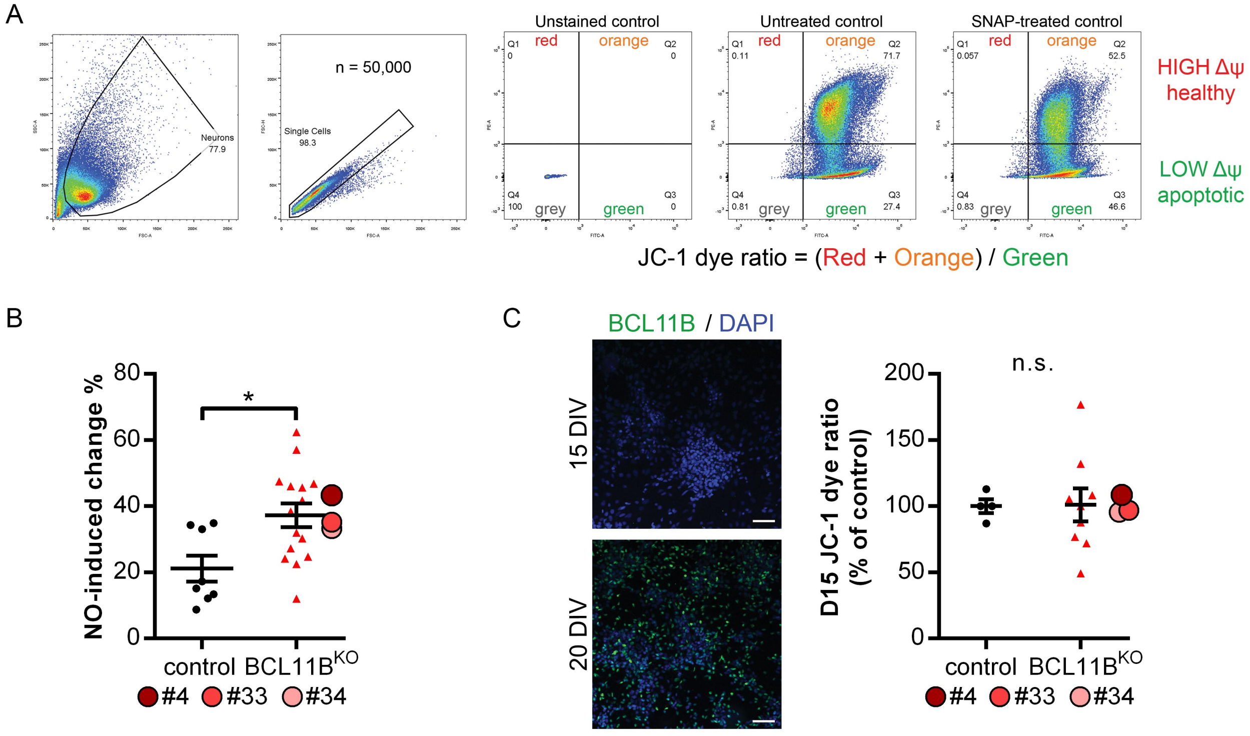


**Figure S2 (Related to Figure 1) – Flow cytometry analysis of mitochondrial metabolic activity.** (**A**) Representative images showing flow cytometry gating strategy to measure mitochondrial membrane potential in ≥50,000 MSNs per sample. The voltage-sensitive dye JC-1 selectively enters mitochondria and changes color from red to green as the membrane potential decreases in response to SNAP-induced oxidative stress. (**B**) Percentage nitric oxide (NO)-induced change in mitochondrial membrane potential compared to untreated condition in MSNs at 40 DIV [Mann-Whitney U test: $U$=25, *$P$=0.023]. (**C**) Images of MSN progenitors in control cultures at 15 (top) and 20 (bottom) DIV immunostained for BCL11B. Scale bars: 50µm. Similar mitochondrial membrane potentials are detected in control and BCL11B$^{KO}$ MSN progenitors at 15 DIV prior to BCL11B expression. All graphs show mean ± s.e.m. for each genotype, with the means for individual clones indicated by red-shaded circles next to BCL11B$^{KO}$ data; n.s., not significant.

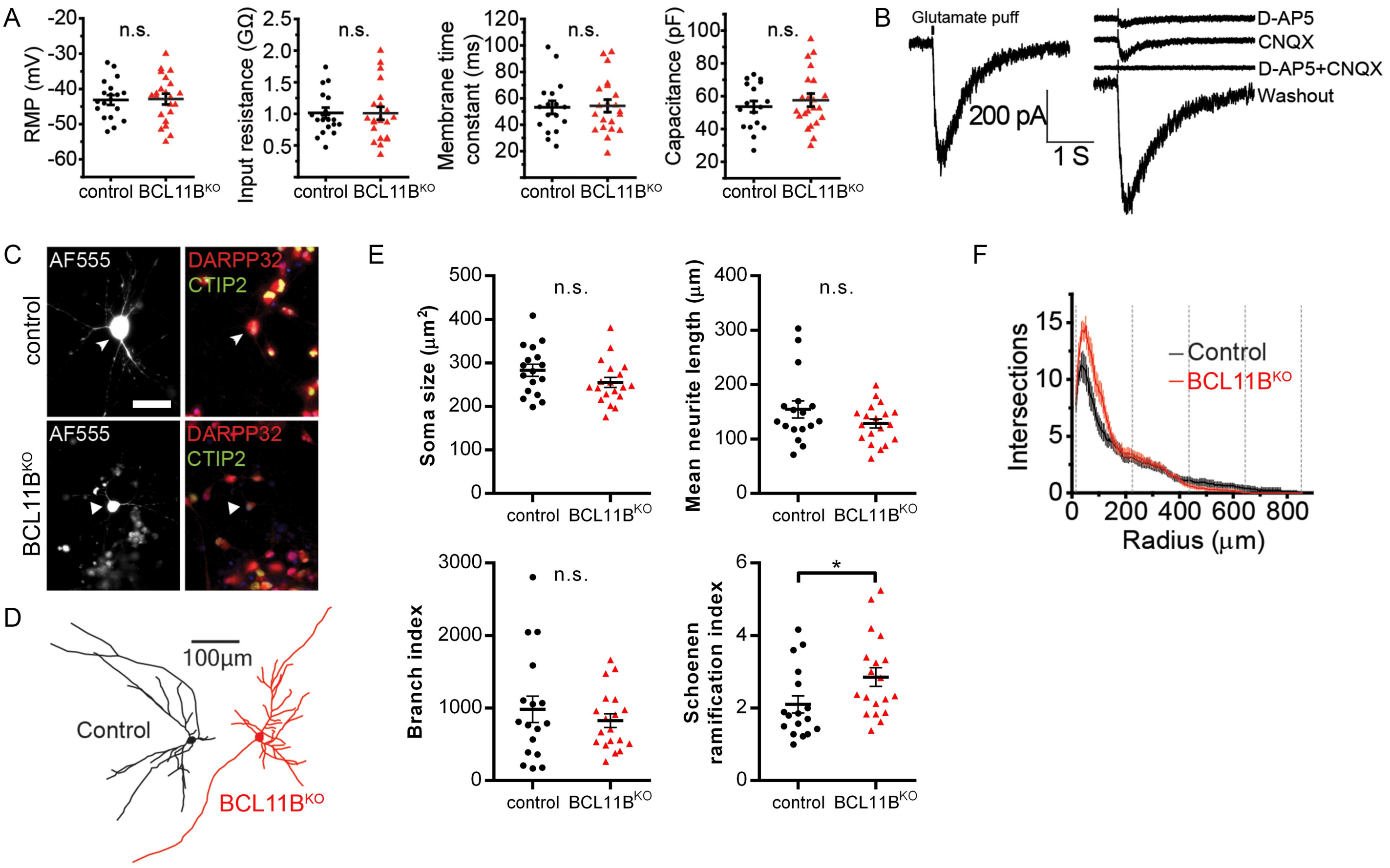


**Figure S3 (Related to Figure 3) – Electrophysiological and morphological analysis of BCL11B$^{KO}$ and control MSNs.** (**A**) All basic membrane properties, including resting membrane potential (RMP), input resistance, membrane time constant and capacitance, were measured and found to not differ significantly between the genotypes. (**B**) Current responses to a DL-glutamic acid puff (200 µM, 30 ms) in control MSNs are mediated by both AMPA and NMDA receptors as indicated by their block by CNQX (50 µM) and D-AP5 (50 µM), respectively. (**C**) Post hoc immunostaining of AF555-filled cells (white, shown by arrow) for BCL11B/CTIP2 and DARPP32 with nuclei counter-stained with DAPI (blue). The BCL11B/CTIP2 antibody binds to exon 1 of the protein, which is upstream of the truncation. Scale bar: 25µm. (**D**) Representative traces of control (black) and BCL11B$^{KO}$ (red) neurons reconstructed in Neurolucida from images of AF555-filled cells. (**E**) Morphological feature analysis in control and BCL11B$^{KO}$ MSNs showing striking similarities between the genotypes including soma size, mean neurite path length, branch index and Schoenen ramification index [Mann-Whitney U test: Schoenen – $U$=88.5, *$P$=0.019]. (**F**) Sholl plots of control and BCL11B$^{KO}$ MSNs with the total area under the curve (AUC) divided into 4 equal segments for statistical analysis [two-way ANOVA: Genotype: $F_{(1,136)}$=1.587, $P$=0.210; AUC: $F_{(3,136)}$=186.063, $P$=6.09x10$^{-48}$]. All line graphs and dot plots depict mean ± s.e.m. for each genotype; n.s., not significant.

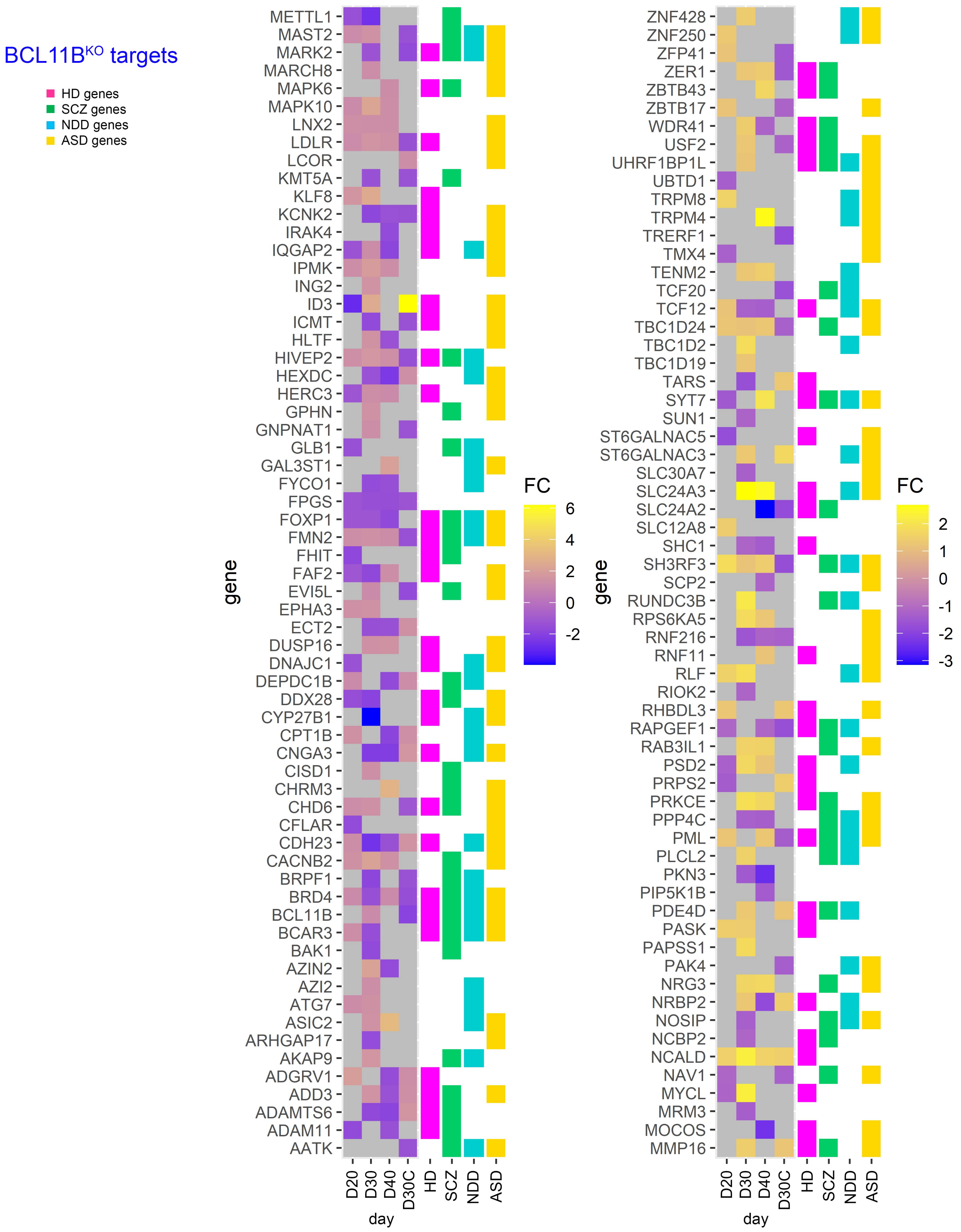


**Figure S4 (Related to Figure 4) – Heatmap of BCL11B target genes within MSN and cortical DEG lists showing their corresponding fold change** (MSN: D20, D30, D40; cortical: D30C). Next to gene name, color icons indicate significant association (differential expression or identified risk variant) with Huntington's disease (HD, pink), schizophrenia (SCZ, green), neurodevelopmental disorder (NDD, blue), and autism spectrum disorder (ASD, yellow).

| Publication | Gene Count | Description |
|---|---|---|
| **HD** | **8886 unique genes** | |
| 2006 Hodges A et al. | 8082 | Transcriptomic analysis of human HD brain |
| 2015 Ring KL et al. | 4454 | RNA-sequencing of human HD neural stem cells |
| 2016 Langfelder P et al. | 3738 | Genomic and proteomic analysis of CAG length-dependent mouse brain |
| 2017 HD iPSC Consortium | 918 | RNA-sequencing of human HD neurons |
| **SCZ** | **4504 unique genes** | |
| 2011 SCZ-PGC | 51 | PGC1 GWAS compilation |
| 2012 Alayew M et al. | 145 | Convergent functional genomics approach to GWAS data |
| 2012 Kirov G et al. | 235 | De novo CNV genes |
| 2013 Ripke S et al. | 114 | GWAS data |
| 2014 Fromer M et al. | 923 | Exome sequencing study of de novo mutations in SCZ |
| 2014 Purcell SM et al. | 1796 | Exome sequencing of SCZ patients |
| 2014 SWG-PGC | 353 | PGC2 GWAS meta-analysis |
| 2018 Pardiñas AF et al. | 795 | CLOZUK and PGC data meta-analysis |
| 2018 Rees E et al. | 187 | Targeted sequencing of SCZ-associated genes |
| 2018 Ruderfer DM et al. | 6 | Genomic dissection of bipolar disorder and SCZ |
| 2018 Schijven D et al. | 6 | Comprehensive pathway analyses of SCZ risk loci |
| 2019 Liu C et al. | 2 | Comprehensive update of findings from candidate gene studies |
| 2019 Rees E et al. | 42 | De novo mutations in SCZ identified by exome sequencing |
| 2019 Lam M et al. | 1194 | Comparison of SCZ GWAS in East Asian and European populations |
| 2019 Gamazon ER et al. | 563 | Multi-tissue transcriptome analyses in psychiatric disorders |
| 2019 Huckins LM et al. | 224 | SCZ risk gene expression across multiple brain regions |
| 2020 Wu Y et al. | 1654 | Validated de novo mutations within genes from denovodb v1.5: SCZ |
| **NDD** | **5561 unique genes** | |
| 2016 McRae JF et al. | 93 | De novo mutations in developmental disorders |
| 2019 Rees E et al. | 22 | De novo mutations in SCZ identified by exome sequencing |
| 2020 Wu Y et al. | 17653 | Validated de novo mutations within genes from denovodb v1.5: NDD |
| **ASD** | **11728 unique genes** | |
| 2014 Fromer M et al. | 1052 | Exome sequencing study of de novo mutations in SCZ |
| 2015 Cotney J et al. | 3051 | Autism-associated chromatin modifier CHD8 list |
| 2020 Wu Y et al. | 116537 | Validated de novo mutations within genes from denovodb v1.5: ASD |

**Figure S5 (Related to Figures 4 and 5)** – Summary of published literature including transcriptomics, GWAS datasets and other functional genomic studies of Huntington's disease (HD), schizophrenia (SCZ), neurodevelopmental disorder (NDD), and autism spectrum disorder (ASD) used to collate neurological disease gene sets for gene enrichment analysis of BCL11B-regulated DEGs. See also **Table S5**.

### Supplemental Table Titles and Legends

**Table S1 (Related to Figure 4) – Differentially expressed genes in $BCL11B^{KO}$ MSNs and cortical neurons.** Gene lists are presented in a separate sheet for each sample (padj<0.01). The first five columns identify the gene, and the other columns contain differential expression statistics for $BCL11B^{KO}$ versus control group comparisons. MSN, medium spiny neuron; CTX, cortical glutamatergic neuron.

**Table S2 (Related to Figure 4) – Summary of KEGG gene set and IPA canonical pathway enrichment analysis of $BCL11B^{KO}$ differentially expressed genes.** This table contains one sheet per sample/analysis combination showing significantly dysregulated pathways (padj<0.05), enrichment statistics and associated gene identifiers. Tables #i-k show pathway analysis results when BCL11B target genes are excluded from MSN30 and CTX30 differentially expressed gene lists. MSN, medium spiny neuron; CTX, cortical glutamatergic neuron.

**Table S3 (Related to Figure 4) – Summary of HD dataset enrichment analysis of the $BCL11B^{KO}$ differentially expressed genes.** The first sheet provides an overview and Fisher's exact test statistics for each association tested. The following four sheets provide a compilation of MSN and cortical differentially expressed genes concordantly dysregulated (padj<0.01) in at least one HD gene expression dataset. Each sheet corresponds to one sample and provides gene identifiers with differential expression statistics. MSN, medium spiny neuron; CTX, cortical glutamatergic neuron.

**Table S4 (Related to Figure 4) – Summary of KEGG gene set and pathway enrichment analysis of concordantly dysregulated genes between $BCL11B^{KO}$ and HD models.** The first sheet provides an overview and Fisher's exact test statistics for HD gene enrichment in core BCL11B-regulted pathways. The following tables contain one sheet per sample showing significantly dysregulated KEGG pathways (padj<0.05), enrichment statistics and associated gene identifiers. MSN, medium spiny neuron; CTX, cortical glutamatergic neuron.

**Table S5 (Related to Figures 5 and S5) – Overview of psychiatric disorder datasets used for enrichment analysis of the BCL11B$^{KO}$ differentially expressed genes.** The first sheet provides a summary of published literature including transcriptomics, GWAS datasets and other functional genomics studies of schizophrenia (SCZ), neurodevelopmental disorder (NDD) and autism spectrum disorder (ASD) used to collate neurological disease gene sets for gene enrichment analysis of BCL11B-regulated differentially expressed genes. Gene lists are presented in a separate sheet for each disorder including gene identifiers and accession information in published literature.

**Table S6 (Related to Figure 5) – Summary of psychiatric disorder dataset enrichment analysis of the BCL11B$^{KO}$ differentially expressed genes.** The first sheet provides an overview and Fisher's exact test statistics for each association tested. The following sheets provide lists of MSN and cortical differentially expressed genes enriched for psychiatric disorder risk variants, each sheet corresponding to one 'sample'/'disease dataset' combination and containing gene identifiers with differential expression statistics. MSN, medium spiny neuron; CTX, cortical glutamatergic neuron.

**Table S7 (Related to Figure 5) – Summary of KEGG gene set and pathway enrichment analysis of BCL11B-regulated genes enriched for psychiatric disorder risk variants.** The first sheet provides an overview and Fisher's exact test statistics for psychiatric risk gene enrichment in core BCL11B-regulted pathways.

The following tables contain one sheet per ‘sample’/‘disease dataset’ combination showing significantly dysregulated KEGG pathways (padj<0.05), enrichment statistics and associated gene identifiers. MSN, medium spiny neuron; CTX, cortical glutamatergic neuron.

### Materials and Methods

#### Immunocytochemistry

Cultured cells were rinsed in PBS and fixed in 3.7% PFA for 15 minutes. Cells were permeabilized in 0.3% Triton-X-100 solution in PBS (PBS-T) and then blocked in PBS-T containing 1% BSA and 3% donkey serum. Cells were incubated with primary antibodies in blocking solution overnight at 4°C. Following three PBS-T washes, Alexa-Fluor secondary antibodies (Thermo Fisher Scientific) were added at 1:1000 in blocking solution for 1 hour at ambient temperature in the dark. Cells were stained with DAPI at 1:1000 (Thermo Fisher Scientific). The following primary antibodies were used for the immunofluorescence studies: rabbit anti-DARPP32 (Santa Cruz Biotechnology #sc-11365, 1:500), rat anti-BCL11B/CTIP2 (Abcam # ab18465, 1:500), rabbit anti-FOXP2 (Abcam #ab16046, 1:500), mouse anti-FOXP1 (Abcam # ab32010, 1:800), rabbit anti-TBR1 (Abcam # ab31940, 1:500), mouse anti-SATB2 (Abcam #ab31940), anti-goat FOXA2 (Santa Cruz Biotechnology #sc-6554), anti-rabbit LMX1A (Sigma #HPA03008), anti-mouse TH (Millipore #MAB318) . Images were taken on a Zeiss LSM710 confocal microscope from at least 5 randomly selected fields/sample and staining quantification was acquired manually in ImageJ (imagej.net) from >5,000 cells/sample blind to the experimental condition.

#### Mitochondrial Assay

The cell-permeant mitochondrial membrane potential sensor JC-1 (Thermo Fisher Scientific) was added directly to live cells at a final concentration of 1 µg/mL for 30 minutes at 37°C. Thus, mitochondrial activity was assessed in control and $BCL11B^{KO}$ cells using flow cytometry with the voltage-sensitive dye JC-1 that selectively accumulates in mitochondria and undergoes a fluorescence emission shift from 590 nm (red) to 525 nm (green) upon membrane potential depolarization allowing the JC-1 dye fluorescence ratio to be used as a relative measure of mitochondrial membrane potential (**Figure S2A**). Where indicated cells were pre-incubated with a nitric oxide donor (S)-Nitroso-N-acetylpenicillamine (SNAP, 1000 µM, Tocris) for 24 hours. All samples plus an unstained control were collected as a single cell suspension dissociated in Accutase (Merck Sigma-Aldrich), centrifuged at 150 g, resuspended in PBS and filtered with a 35 µm nylon mesh. Samples were analyzed on a flow cytometer according to the manufacturer's protocol ($488_{Ex}$, FL1-$525_{Em}$ and FL2-$590_{Em}$). Measurements were taken from >50,000 single cells after excluding debris and selecting singlets (**Figure S2A**). Mitochondrial depolarization was measured as a decrease in the (red+orange)/green fluorescence intensity ratio.

**Live-cell Calcium Imaging**

Neuronal cultures were cultured in 24-well plates and incubated with a $Ca^{2+}$-sensitive probe Fluo-4-AM (Thermo Fisher Scientific) at a final concentration of 5 µM in N2B27 medium for 30 minutes at 37 °C. The solution was then aspirated and replaced with pre-warmed artificial cerebrospinal fluid (aCSF) solution [2mM $CaCl_2$, 142 mM NaCl, 2.5mM KCl, 1mM $MgCl_2$, 10mM HEPES, 30mM D-glucose – all from Merck Sigma-Aldrich]. Plates were then transferred to a heated environmental chamber atop an inverted Zeiss AG Axio Observer D1 inverted microscope stage and allowed to equilibrate for 10 minutes.

For thapsigargin (Tg, Tocris) treatment paradigm, $Ca^{2+}$-free aCSF was supplemented with 0.16 mM EGTA (Merck Sigma-Aldrich) to chelate extracellular $Ca^{2+}$. Cultures were first incubated in 1 µM Tg in $Ca^{2+}$-free aCSF to deplete neurons of intracellular $Ca^{2+}$ before switching the solution to 1 µM Tg in the standard 2mM $Ca^{2+}$ aCSF (**Figure 1E**). KCl was used at 130 mM in the standard aCSF solution

(**Figure 1F**). To modulate PKA/DRD1-mediated $Ca^{2+}$ signaling in neurons, 10 µM Roscovitine, 10 µM SKF-81297 and 10 µM 8-bromo-cAMP (all from Tocris) solutions in the standard aCSF were prepared fresh on the day of recording. Spontaneous activity- and drug-evoked $Ca^{2+}$ transients were recorded over 10 minutes (**Figure 2C**). DL-Glutamic acid (Merck Sigma-Aldrich) was dissolved in aCSF at a final concentration of 100 µM and applied to neurons in 1-minute pulses followed by 1-minute washout interval over a 40-minute recording window (**Figure 3D**). All drug solutions were pre-warmed to 37°C and delivered/aspirated to/from cultured cells via a pump delivery system.

To visualize intracellular $Ca^{2+}$, cultures were excited at 488 nm and fluorescence emission was detected every 100 ms while the camera shutter was kept open for the duration of the recording. Subsequent data processing was performed using FluoroSNNAP algorithm according to the protocol (1). Individual traces were normalized to initial fluorescence ($F/F_0$) and averaged across samples. The line graphs show mean fluorescence (black/red thick lines) ± s.e.m. (grey/pink thin lines) plotted versus time. The mean maximum change in Fluo-4-AM fluorescence (Max $\Delta F/F_0$) was determined by averaging the peak $F/F_0$ calculated for each trace of a given sample. Responses to glutamate pulses were quantified manually by inspecting individual traces and then averaged across samples.

**Electrophysiology**

*Astrocyte isolation and culture*

The cortex of postnatal day 7 C57BL/6 mouse pups was dissected and dissociated in 2.5% trypsin. Cortical cell suspensions were plated in PDL-coated T75 flasks and grown in DMEM/F12 with 10% fetal bovine serum (Thermo Fisher Scientific), Lglutamine and MycoZap Plus-CL (Lonza). Once confluent, flasks were shaken vigorously for 30 minutes to remove microglia. Remaining astrocytes were then dissociated with 2.5% trypsin and seeded onto 13 mm glass coverslips co-coated with PDL and laminin.

*Electrophysiology cell preparation*

At 20 days of differentiation (DIV), medium spiny neuron (MSN) progenitors were resuspended in Matrigel [1:15 diluted in N2B27 with ROCK inhibitor (10 μM, Tocris), activin A and BDNF] and plated in droplets onto PDL and laminin co-coated 6 mm glass coverslips in a 30 mm petri dish (2, 3). After 90 minutes, wells were flooded with medium (N2B27 supplemented with ROCK inhibitor, activin A and BDNF), and one astrocyte coverslip was placed in each dish for medium conditioning. The following day, medium was changed to BrainPhys basal medium (STEMCELL Technologies) supplemented with B27, activin A and SCM1 factors – BDNF (10 ng/ml), LM22A4 (1 μM, Merck Sigma-Aldrich), DAPT (10 μM, Tocris), PD0332991 (2 μM, Merck Sigma-Aldrich), CHIR99021 (3 μM, Tocris), Forskolin (10 μM, Tocris), GABA (300 μM, Tocris), ascorbic acid (200 μM, Merck Sigma-Aldrich) and $CaCl_2$ (0.5 mM) (4). SCM1 factors were all removed at around 30 DIV except BDNF and ascorbic acid.

*Electrophysiology methods*

Whole-cell patch-clamp recordings were acquired from control and $BCL11B^{KO}$ #4 MSNs at 40 DIV at room temperature using a MultiClamp 700B amplifier and Digidata 1550 digitizer with pCLAMP 10 software (Molecular Devices). For current-clamp recordings, aCSF contained (in mM): 135 NaCl, 5 KCl, 1.2 $MgCl_2$, 1.25 $CaCl_2$, 10 D-glucose, 5 HEPES, pH 7.4. Recording pipettes (3-8 MΩ) were pulled from borosilicate capillary glass (Sutter Instruments, USA) and filled with internal solution containing (in mM): 117 KCl, 10 NaCl, 11 HEPES, 2 $Na_2$-ATP, 2 Na-GTP, 1.2 $Na_2$-phosphocreatine, 2 $MgCl_2$, 1 $CaCl_2$ and 11 EGTA (all from Merck Sigma-Aldrich), pH 7.2. Resting membrane potential was averaged from 10 seconds of recording immediately after break-in. Input resistance ($R_N$) was calculated using Ohm's law from the amplitude of the average steady-state voltage deflections in response to a series of 1-second long hyperpolarizing current injection steps (-10 pA). Membrane capacitance ($C_m$) was calculated according to $C_m = \tau_m/R_N$, where the membrane time-constant ($\tau_m$) was derived from a single exponential function fit to the repolarizing phase of the voltage response evoked by hyperpolarizing current injection. For voltage-clamp recordings, aCSF composition was the same as above except $MgCl_2$ was omitted and glycine (1 µM, Merck Sigma-Aldrich) was added. Internal pipette solution was the same as above except KCl was substituted with 117 mM K-Gluconate (Merck Sigma-Aldrich). SKF81297 (10 µM) was added to aCSF where specified. Cells were held at -70 mV and series resistance was monitored using regular -10 mV steps. A second glass pipette was filled with aCSF containing DL-glutamate (200 µM), which was focally applied (30 ms) to the patched cell and the evoked current response recorded. Data were analyzed with Clampfit software (Molecular Devices) then exported to and plotted using Origin (OriginLab).

Alexa Fluor hydrazide 555 (Thermo Fisher Scientific) was included in the internal solution (1 mM) for morphological analysis. At the end of recording, AF555 dye-filled cells were imaged *in situ* for morphological analysis. Imaged cells were reconstructed and Sholl analysis carried out in Neurolucida (MBF Bioscience) and data was exported to Origin for analysis. The area under the curve was divided into 4 equal segments to detect differences in neurite branching and calculated from the number of intersections at each radius. The Schoenen ramification index was calculated as the ratio between maximum number of intersections and number of primary neurites.

**RNA Sequencing Data Analysis**

Total RNA was extracted from TRIzol lysates using the PureLink RNA mini kit (Ambion) from three biological replicates per genotype at 20, 30 and 40 DIV for MSNs and 30 DIV for cortical glutamatergic neurons. These time points were chosen to reflect the onset of BCL11B expression in nascent post mitotic neurons and subsequent rapid increase in BCL11B levels during differentiation, as well as to correspond to the time points at which several cellular pathologies have been observed. Paired-end sequencing was performed at Oxford Genomic Centre on an Illumina HiSeq 4000 (Illumina, San Diego, USA) obtaining a library size of ~80 million reads per sample. RNA-seq data reported in this paper are available with the SRA accession numbers **PRJNA474679** and **PRJNA767962** (https://www.ncbi.nlm.nih.gov/sra/PRJNA474679 and https://www.ncbi.nlm.nih.gov/sra/PRJNA767962). FASTQ files were trimmed and mapped to the Ensembl human genome GRCh38.84 (hg38) using STAR (v2.5.1b). Quality of the samples was assessed using FastQC (v0.11.2) prior to and after trimming. Gene counts were obtained from the number of uniquely aligned unambiguous reads by Subread:featureCount (v1.4.6-p2). Library size normalization and differentially expressed genes (DEGs) were determined using the R/Bioconductor package DESeq2 (v1.14.1) (5). Benjamini-Hochberg (BH) test for multiple correction was used to control the false discovery rate (FDR). Subsequent study of KEGG gene set and pathway enrichment analysis was performed on protein coding DEGs with an EntrezID and FDR<0.01 using the Ingenuity Pathway Analysis (Qiagen) and R/Bioconductor package clusterProfiler (v3.12.14) (6). The associations between present BCL11B$^{KO}$ gene sets and published Huntington's disease, schizophrenia, neurodevelopmental disorder and autism spectrum disorder gene sets were determined by Fisher's exact test, performed in RStudio (www.r-project.org).

**Statistical Analysis**

RNA sequencing data was analyzed separately as described above. SPSS Statistics 25 software (IBM) was used for all other statistical analyses except patch-clamp data, which was analyzed in Origin. All quantified data were plotted in Prism 6 (GraphPad Software) and are reported as mean ± s.e.m. with statistical details presented in the figure legends. For normally distributed data as determined by the Shapiro-Wilk test, we performed either two-tailed Student's *t*-test or two-way ANOVAs followed by post hoc Bonferroni correction for multiple comparisons. When data was not modeled by a normal distribution, analysis was subjected to nonparametric Mann-Whitney U test or Kruskal-Wallis test followed by post hoc Bonferroni test. Differences in sample distributions were considered statistically significant at $p<0.05$. No statistical methods were used to predetermine sample sizes, but our sample sizes are similar to those reported in previous publications (2, 7, 8). Randomization was used to assign experimental conditions and collect data, and data collection was always done in parallel with controls. Data analyses were not performed blind to the conditions of the experiments except where stated otherwise.

1. Patel TP, Man K, Firestein BL, Meaney DF (2015): Automated quantification of neuronal networks and single-cell calcium dynamics using calcium imaging. *J Neurosci Methods*. 243:26-38.
2. Arber C, Precious SV, Cambray S, Risner-Janiczek JR, Kelly C, Noakes Z, et al. (2015): Activin A directs striatal projection neuron differentiation of human pluripotent stem cells. *Development*. 142:1375-1386.
3. Fjodorova M, Noakes Z, Li M (2020): A role for TGFβ signalling in medium spiny neuron differentiation of human pluripotent stem cells. *Neuronal Signaling*. 4:NS20200004.

4. Telezhkin V, Schnell C, Yarova P, Yung S, Cope E, Hughes A, et al. (2016): Forced cell cycle exit and modulation of GABA(A), CREB, and GSK3 beta signaling promote functional maturation of induced pluripotent stem cell-derived neurons. *American Journal of Physiology-Cell Physiology*. 310:C520-C541.
5. Love MI, Huber W, Anders S (2014): Moderated estimation of fold change and dispersion for RNA-seq data with DESeq2. *Genome Biol*. 15.
6. Yu G, Wang L-G, Han Y, He Q-Y (2012): clusterProfiler: an R Package for Comparing Biological Themes Among Gene Clusters. *Omics-a Journal of Integrative Biology*. 16:284-287.
7. Victor MB, Richner M, Olsen HE, Lee SW, Monteys AM, Ma C, et al. (2018): Striatal neurons directly converted from Huntington's disease patient fibroblasts recapitulate age-associated disease phenotypes. *Nat Neurosci*. 21:341-352.
8. Cavey M, Collins B, Bertet C, Blau J (2016): Circadian rhythms in neuronal activity propagate through output circuits. *Nat Neurosci*. 19:587-595.
